# Complete map of SARS-CoV-2 RBD mutations that escape the monoclonal antibody LY-CoV555 and its cocktail with LY-CoV016

**DOI:** 10.1101/2021.02.17.431683

**Authors:** Tyler N. Starr, Allison J. Greaney, Adam S. Dingens, Jesse D. Bloom

**Affiliations:** Basic Sciences and Computational Biology, Fred Hutchinson Cancer Research Center, Seattle, WA 98109; Department of Genome Sciences, University of Washington, Seattle, WA 98109; Medical Scientist Training Program, University of Washington, Seattle, WA 98109; Howard Hughes Medical Institute, Seattle, WA 98109

## Abstract

Monoclonal antibodies and antibody cocktails are a promising therapeutic and prophylaxis for COVID-19. However, ongoing evolution of SARS-CoV-2 can render monoclonal antibodies ineffective. Here we completely map all mutations to the SARS-CoV-2 spike receptor binding domain (RBD) that escape binding by a leading monoclonal antibody, LY-CoV555, and its cocktail combination with LY-CoV016. Individual mutations that escape binding by each antibody are combined in the circulating B.1.351 and P.1 SARS-CoV-2 lineages (E484K escapes LY-CoV555, K417N/T escape LY-CoV016). Additionally, the L452R mutation in the B.1.429 lineage escapes LY-CoV555. Furthermore, we identify single amino acid changes that escape the combined LY-CoV555+LY-CoV016 cocktail. We suggest that future efforts should diversify the epitopes targeted by antibodies and antibody cocktails to make them more resilient to antigenic evolution of SARS-CoV-2.

## Introduction

Monoclonal antibodies have been rapidly developed for the treatment and prophylaxis for COVID-19 where they have shown promise in humans [1,2] and animal models [3–7]. One leading antibody is LY-CoV555 (bamlanivimab) [4], which has an emergency use authorization (EUA) for the therapeutic treatment of COVID-19 [8]. An EUA was also recently granted for administration of LY-CoV555 as a cocktail with another antibody, LY-CoV016 (also known as etesevimab) [9].

A key question is whether SARS-CoV-2’s ongoing evolution will lead to escape from these antibodies. This question has taken on growing importance with the recent emergence of SARS-CoV-2 lineages containing mutations in the spike receptor-binding domain (RBD) [10–13], the target of the most clinically advanced antibodies including LY-CoV555 and LY-CoV016. A flurry of recent studies have addressed this question by characterizing the antigenic impacts of the mutations in these emerging lineages—and unfortunately, some of the mutations in emerging lineages reduce binding and neutralization by some key antibodies in clinical development, including LY-CoV555 and LY-CoV016 [14–17].

To enable more comprehensive and prospective assessment of the impacts of viral mutations, we recently developed a method to completely map how all single amino-acid mutations in the SARS-CoV-2 RBD affect antibody binding [15,18,19]. These maps enable immediate interpretation of the consequences of new mutations and systematic comparison of escape mutations across antibodies.

Here, we prospectively map how all mutations to the RBD affect binding by LY-CoV555 alone and in a cocktail with LY-CoV016. (We have previously reported how all mutations affect binding by LY-CoV016 alone [15]). Binding by LY-CoV555 is escaped by mutations within and near the RBD “receptor-binding ridge”, including by mutations at sites L452 and E484 that are present in emerging viral lineages. Furthermore, the LY-CoV555+LY-CoV016 cocktail is escaped by the specific combinations of mutations at K417 and E484 found in the B.1.351 and P.1 lineages. Finally, we show that several individual amino-acid mutations are capable of escaping the combined LY-CoV555+LY-CoV016 cocktail.

## Results

We applied a previously described deep mutational scanning approach to comprehensively map mutations in the SARS-CoV-2 RBD that escape binding from antibodies [15,18,19]. Briefly, this method involves displaying nearly all amino-acid mutants of the SARS-CoV-RBD on the surface of yeast [20], incubating the yeast with an antibody or antibody cocktail, using fluorescence-activated cell sorting (FACS) to enrich functional RBD mutants that escape antibody binding (Fig. S1), and using deep sequencing to quantify the extent to which each mutation is enriched in the antibody-escape population relative to the original population. The effect of each mutation is quantified by calculating its “escape fraction,” which represents the fraction of yeast expressing this mutant that fall in the antibody-escape FACS bin (these fractions range from 0 for mutations with no effects to 1 for mutations that strongly escape antibody binding).

We used this approach to map how all RBD mutations affect binding by a recombinant form of LY-CoV555 and its 1:1 cocktail combination with recombinant LY-CoV016, and examined these maps alongside similar data [15] that we recently reported for LY-CoV016 alone (Figures 1A, S1; interactive visualizations at https://jbloomlab.github.io/SARS-CoV-2-RBD_MAP_LY-CoV555/). The maps show that LY-CoV555 is escaped by mutations at a focused set of sites, with site E484 standing out as a hotspot of escape (Figure 1A). We layered onto the escape maps our previous deep mutational scanning measurements [20] of how mutations affect ACE2 binding (Fig. 1A) or expression of folded RBD (Fig. S2), and found that mutations escaping LY-CoV555 often have no adverse effect on these two functional properties of the RBD.

**Figure 1.**
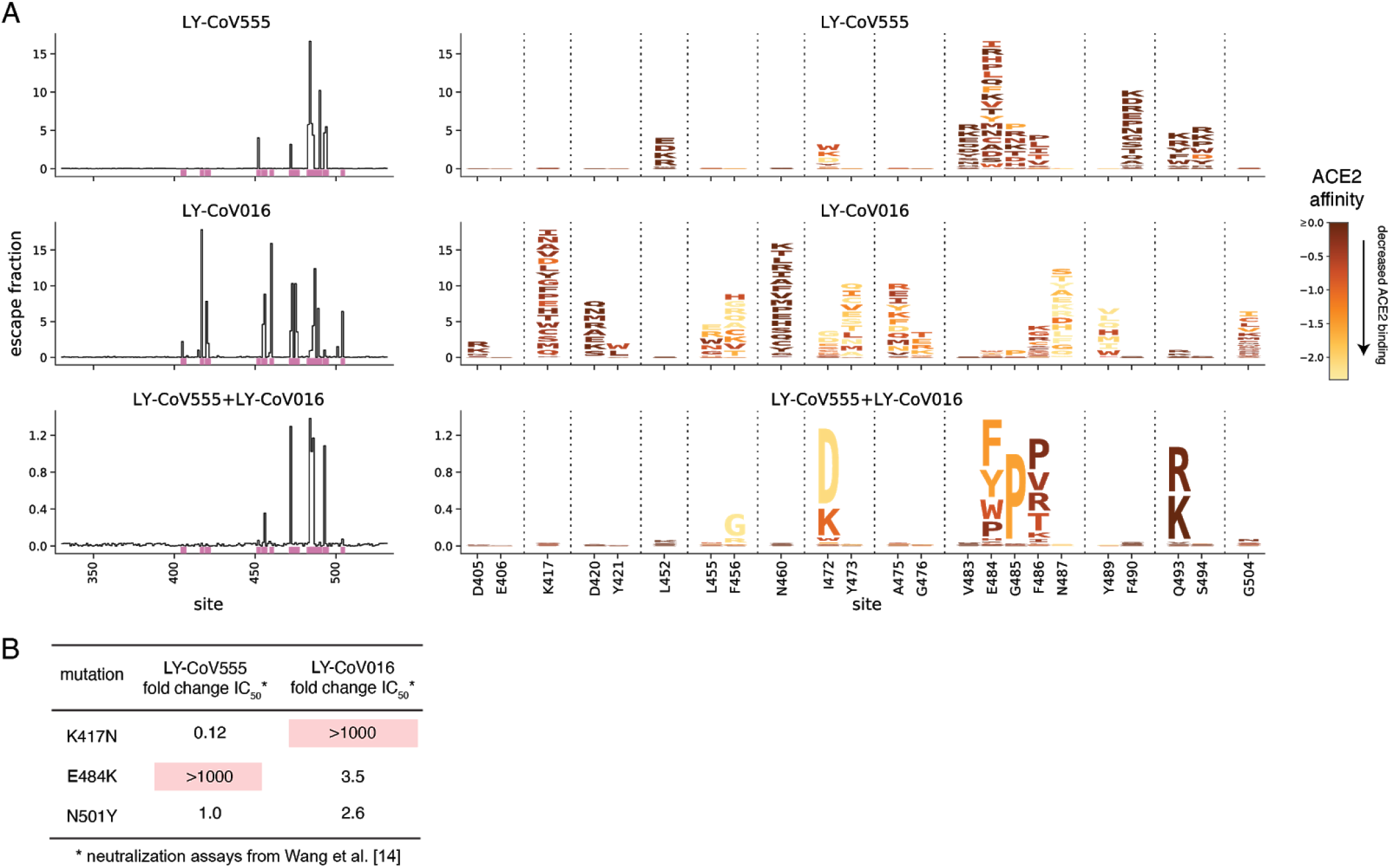
Comprehensive escape maps for LY-CoV555, LY-CoV016, and a 1:1 cocktail of the two antibodies. (A) Newly described escape maps for LY-CoV555 and LY-CoV555+LY-CoV016 cocktail, alongside our previously reported escape map for LY-CoV016 [15]. Line plots at left show the total escape (sum of per-mutation escape fractions) at each RBD site. Sites indicated by pink lines on the x-axis are then shown in zoomed in form in the logoplots at right. In these logoplots, the height of each letter indicates the escape fraction for that mutation (larger letters mean stronger escape from antibody binding). Letters are colored by how mutations impact ACE2 binding affinity (scale bar bottom right), as measured in our prior deep mutational scan [20]. See Fig. S2 for escape maps colored by mutation effects on folded RBD expression. Note that y-axis is scaled differently for each antibody / cocktail. The sites shown in the logoplots are those where mutations have an appreciable effect on either antibody, as well as site 406 (which is an escape mutation from the REGN-COV2 cocktail [15]). (B) Literature measurements of the effects of K417N, E484K, and N501Y on neutralization by LY-CoV555 and LY-CoV016 [14]. These measurements validate our maps, which suggest that K417N specifically escapes LY-CoV016, E484K specifically escapes LY-CoV555, and N501Y impacts neither antibody.

Comparison of the LY-CoV555 escape map with a map we previously reported for LY-CoV016 shows that the latter antibody is primarily escaped by mutations at sites where mutations do not affect LY-CoV555 (e.g., K417 and N460; Fig 1A, S1). However, there are some sites where single mutations escape binding by both LY-CoV555 and LY-CoV016, and as a result a 1:1 cocktail of the two antibodies is escaped by several single mutations including I472D, G485P, and Q493R/K (Fig. 1A, S2; see the zoomable interactive maps https://jbloomlab.github.io/SARS-CoV-2-RBD_MAP_LY-CoV555/ to examine these mutations at higher resolution). Note that some of the other smaller cocktail escape mutations in the cocktail maps may reflect a higher potency of LY-CoV555 in the 1:1 cocktail rather than representing mutations that truly escape binding by both antibodies. Mutations at position Q493 are notably well tolerated with respect to ACE2 binding and RBD expression (Fig 1A, S2)—indeed, Q493K has been observed in a persistently infected immunocompromised patient [15,21].

The binding measurements in our maps are consistent with previously reported effects of mutations on antibody neutralization from the literature (Fig. 1B). Specifically, Wang et al. [14] have reported that E484K and K417N dramatically and specifically reduce neutralization by LY-CoV555 and LY-CoV016, respectively, while N501Y has no impact on neutralization by either antibody. However, our maps greatly extend this prior knowledge by identifying all mutations at all positions that impact binding by these antibodies and their combination.

We used the maps to assess how all RBD mutations present in sequenced SARS-CoV-2 isolates impact binding by each antibody (Fig. 2A). The escape mutations present at the highest frequency among sequenced isolates are E484K, L452R, and S494P for LY-CoV555, and K417N/T for LY-CoV016. An array of other mutations that escape each antibody are present at lower frequency among sequenced isolates. Of particular note, the B.1.351 (a.k.a. 20H/501Y.V2) [10] and P.1 (a.k.a. 20J/501Y.V3) [12] lineages contain combinations of mutations (E484K and K417N/T) that individually escape each antibody (Fig. 2B), suggesting that the LY-CoV555+LY-CoV016 cocktail may be ineffective against these lineages. In addition, the B.1.429 lineage (a.k.a. 20C/CAL.20C) that has risen to high frequency in southern California contains L452R [13], which escapes LY-CoV555 (Fig. 2B). We also note that single mutations that escape both antibodies (Q493R and Q493K) have been observed in a handful of sequenced isolates (Fig. 2A).

**Figure 2.**
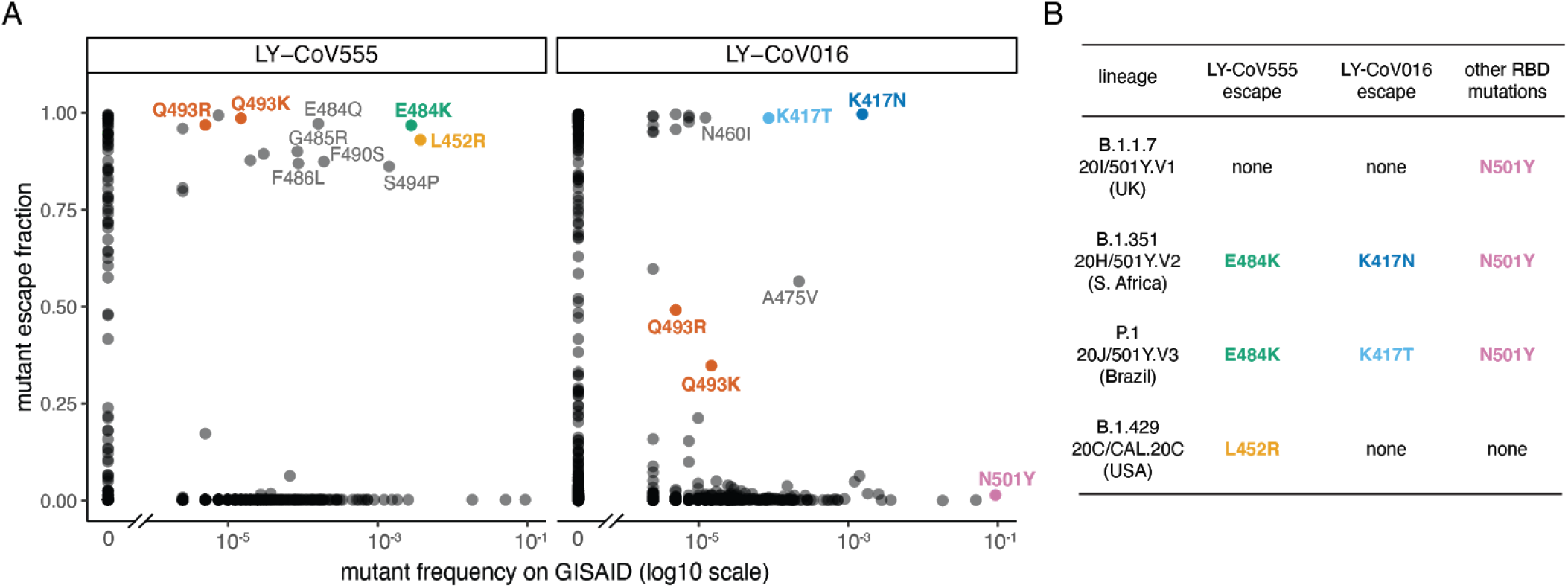
Mutations present in sequenced SARS-CoV-2 isolates that escape antibody binding. (A) For each mutation, the escape fraction measured in the current (LY-CoV555) or prior (LY-CoV016 [15]) study is plotted against the mutation’s frequency among all 405,692 high-quality human-derived SARS-CoV-2 sequences in GISAID as of February 6, 2021. Mutations with notable frequencies are labeled, and those discussed in the text are colored to key with panel (B) or to highlight observed cocktail escape mutations (Q493K/R). (B) The RBD mutations in four emerging viral lineages, categorized by their effect on binding by LY-CoV555 and LY-CoV016. The B.1.351 and P.1 lineages contain combinations of mutations that escape each component of the LY-CoV555+LY-CoV016 cocktail. Lineages described in the following references: B.1.1.7, [11]; B.1.351, [10]; P.1, [12]; B.1.429, [13].

To gain insight into the structural basis for the escape mutations, we projected our escape maps onto crystal structures of the antibodies bound to the RBD [4,22] (Fig. 3, interactive visualizations at https://jbloomlab.github.io/SARS-CoV-2-RBD_MAP_LY-CoV555/). LY-CoV016 and LY-CoV555 bind opposite sides of the “receptor-binding ridge”, a structurally [23] and evolutionarily [24,25] dynamic region of the RBD that forms part of the ACE2 receptor contact surface. The hotspots of escape for each antibody map closely to the core of each antibody-RBD complex. The sites where mutations escape the LY-CoV555+LY-CoV016 cocktail highlight their joint recognition of the receptor-binding ridge (Fig. 3). The cocktail escape site Q493 is not in the receptor-binding ridge, but is in a region of joint structural overlap by the two antibodies, such that the introduction of bulky, positively charged residues (R, K) may directly impact binding by each antibody.

**Figure 3.**
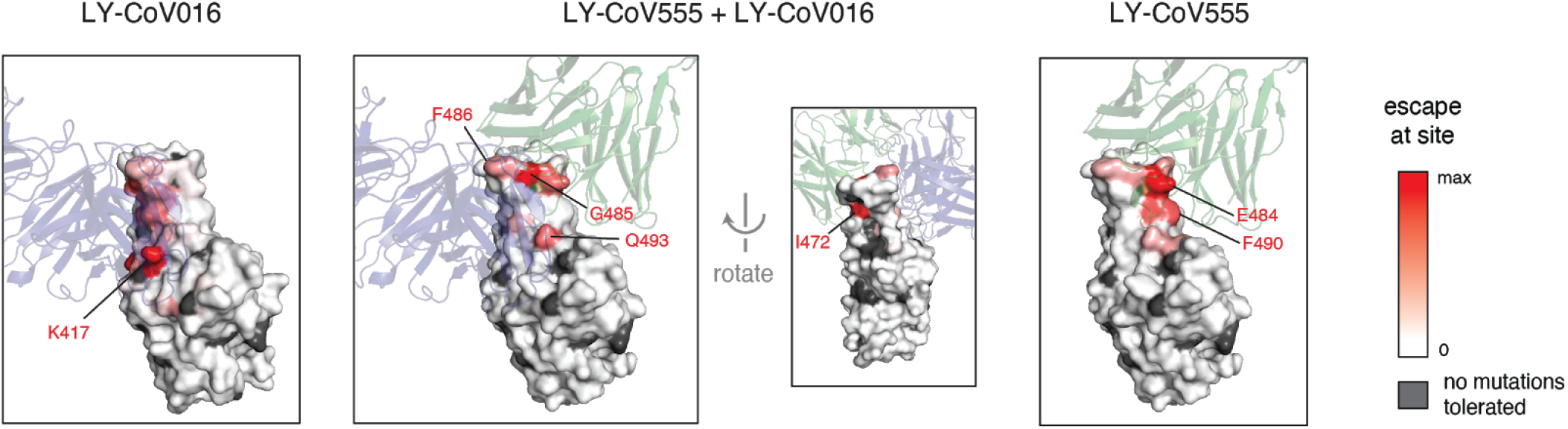
Escape maps projected onto structures of the RBD bound by LY-CoV555 or LY-CoV016. In each structure, the RBD surface is colored by escape at each site (white = no escape, red = strongest site-total escape for antibodies or strongest per-mutation escape for cocktail, gray = no escape because no mutations functionally tolerated). Sites of interest in each structure are labeled. The structures are as follows: LY-CoV016 (PDB 7C01 [22]); LY-CoV555 (PDB 7KMG [4]); cocktail escape projected onto the 7KMG structure, with the LY-CoV016 Fab chain aligned from the 7C01 structure for reference.

## Discussion

We generated complete maps of mutations that escape a leading antibody and antibody cocktail being used to combat COVID-19. Our maps highlight the need to consider circulating SARS-CoV-2 diversity in regions where these antibodies are deployed, as several viral lineages already have mutations that escape binding from LY-CoV555 and its cocktail with LY-CoV016. More broadly, the maps we report will continue to enable immediate assessment of the impacts of newly observed mutants on these antibodies and their cocktail—although it will of course remain necessary to validate key findings with additional virological experiments.

More broadly, our maps suggest that it may be advisable to more systematically consider possible escape mutations when devising antibodies for clinical use against SARS-CoV-2. It is now clear that human coronaviruses undergo antigenic evolution in response to immune pressure [26,27], and we and others have begun to map out the key sites in the RBD that are targeted by human antibody immunity [19,28–30]. The recent rise in frequency of mutations at site E484 suggests that this immunity may be beginning to drive antigenic variation within immunodominant positions in the RBD. Unfortunately, many of the leading therapeutic antibodies target the same epitopes as polyclonal antibody immunity, such as residue E484 or the 443-450 loop [19]. Because the clinical usage of monoclonal antibodies is unlikely to be so widespread as to drive viral evolution in the same way as infection- or vaccine-induced immunity, development of antibodies targeting less immunodominant epitopes might prove to be a strategy that is more resilient to the evolution of SARS-CoV-2.

## Acknowledgements

We thank Bryan Jones and Eli Lilly & Co. for sharing the LY-CoV555 sequence and structure, the Fred Hutch Flow Cytometry and Genomics facilities, and Fred Hutch Scientific Computing, supported by ORIP grant S10OD028685. This was supported by the NIAID (R01AI127893 and R01AI141707 to J.D.B. and T32AI083203 to A.J.G.), and the Gates Foundation (INV-004949 to J.D.B.). T.N.S. is an HHMI Fellow of the Damon Runyon Cancer Research Foundation (DRG-2381-19). J.D.B. is an Investigator of the Howard Hughes Medical Institute.

## Author contributions

All authors designed the study. T.N.S. performed the experiments and analyzed the data. T.N.S. and J.D.B. wrote the initial draft, and all authors edited the final version.

## Competing interests

The authors declare no competing interests.

## Materials and Methods

### Data and Code Availability

- Complete computational pipeline: https://github.com/jbloomlab/SARS-CoV-2-RBD_MAP_LY-CoV555
- Markdown summaries of computational analysis: https://github.com/jbloomlab/SARS-CoV-2-RBD_MAP_LY-CoV555/blob/main/results/summary/summary.md
- Raw data table of mutant escape fractions: Data S1 and https://github.com/jbloomlab/SARS-CoV-2-RBD_MAP_LY-CoV555/blob/main/results/supp_data/LY_cocktail_raw_data.csv
- Raw Illumina sequencing data: NCBI SRA, BioProject: PRJNA639956, BioSample SAMN17836431

### Antibodies

The LY-CoV555 antibody variable domain sequences were acquired from the LY-CoV555 crystal structure file (PDB 7KMG, [4]), which was generously shared by Bryan Jones and Eli Lilly and Co. prior to its publication. Purified antibody was produced by Genscript as human IgG in HD 293F mammalian cells, and affinity purified over RoboColumn Eshmuno A 0.6mL columns. LY-CoV016 was previously produced via the same approach as described in Starr et al. [15].

### Comprehensive profiling of mutations that escape antibody binding

Antibody escape mapping experiments were performed in biological duplicate using a deep mutational scanning approach. Assays were performed exactly as described by Starr et al. [15], based on the approach first described in Greaney et al. [18]. Briefly, yeast-surface display libraries expressing 3,804 of the 3,819 possible amino acid mutations in the SARS-CoV-2 RBD (Wuhan-Hu-1 sequence, Genbank MN908947, residues N331-T531) were previously sorted to select mutants capable of binding human ACE2. Libraries were induced for RBD surface expression and labeled with 400 ng/mL antibody (LY-CoV555, or 200 ng/mL each of LY-CoV555 and LY-CoV016 for 400 ng/mL total antibody). Cells were then incubated with 1:200 PE-conjugated goat anti-human-IgG (Jackson ImmunoResearch 109-115-098) to label for bound antibody and 1:100 FITC-conjugated anti-Myc (Immunology Consultants Lab, CYMC-45F) to label for RBD surface expression. Yeast expressing the unmutated SARS-CoV-2 RBD were prepared in parallel to library samples and labeled at 400 ng/mL and 4 ng/mL with the corresponding antibody/cocktail for setting selection gates.

Antibody-escape cells were selected via fluorescence-activated cell sorting (FACS) on a BD FACSAria II. FACS selection gates (Fig. S1) were drawn to capture 95% of unmutated yeast labeled at the 100x reduced 4 ng/mL antibody labeling concentration. For each sample, 10 million RBD+ cells were processed on the cytometer to sort out antibody-escape cells (fractions shown in Fig. S1B), which were grown out overnight. Plasmid was purified from pre-sort and antibody-escape populations, and mutant frequencies pre- and post-sort were determined by Illumina sequencing of variant-identifier barcodes, exactly as described in Starr et al. [20].

Escape fractions were computed as described in Starr et al. [15]. Briefly, we used the `dms_variants` package (https://jbloomlab.github.io/dms_variants/, version 0.8.2) to process Illumina sequences into counts of each barcoded RBD variant using the barcode/RBD look-up table from Starr et al. [20]. The escape fraction of each library variant was determined as the fraction of cells carrying a particular barcode that were sorted into the antibody-escape bin, using the equation given in Greaney et al. [18]. Scores were filtered for minimum library representation and mutant functionality as described in Starr et al. [15], and single-mutant escape scores were deconvolved using global epistasis models [31]. Markdown summaries of all steps of computational analysis are available on GitHub: https://github.com/jbloomlab/SARS-CoV-2-RBD_MAP_LY-CoV555/blob/main/results/summary/summary.md.

### Circulating variants

All spike sequences present on GISAID [32] as of February 6, 2021 were downloaded and aligned via `mafft` [33]. Sequences from non-human origins, sequences with gaps or ambiguous characters, and sequences with more than 8 RBD mutations from consensus were removed. RBD amino acid differences were enumerated compared to the Wuhan-Hu-1 RBD sequence. We acknowledge all contributors to the GISAID EpiCoV database for their sharing of sequence data. (All contributors listed at: https://github.com/jbloomlab/SARS-CoV-2-RBD_MAP_LY-CoV555/blob/main/data/gisaid_hcov-19_acknowledgement_table_2021_02_06.pdf).

### Data visualization

Static logoplots were created using `dmslogo` (https://jbloomlab.github.io/dmslogo/). Interactive visualizations of the escape maps and their projection onto the ACE2-bound (PDB: 6M0J, [34]) and antibody-bound structures available at https://jbloomlab.github.io/SARS-CoV-2-RBD_MAP_LY-CoV555/ were created using `dms-view` (https://dms-view.github.io/docs/, [35]). For Fig. 3, escape scores were mapped to PDB b-factors and visualized in PyMol using antibody-bound RBD structures PDB 7KMG [4] and PDB 7C01 [22].

---

**Supplemental Figure 1.**
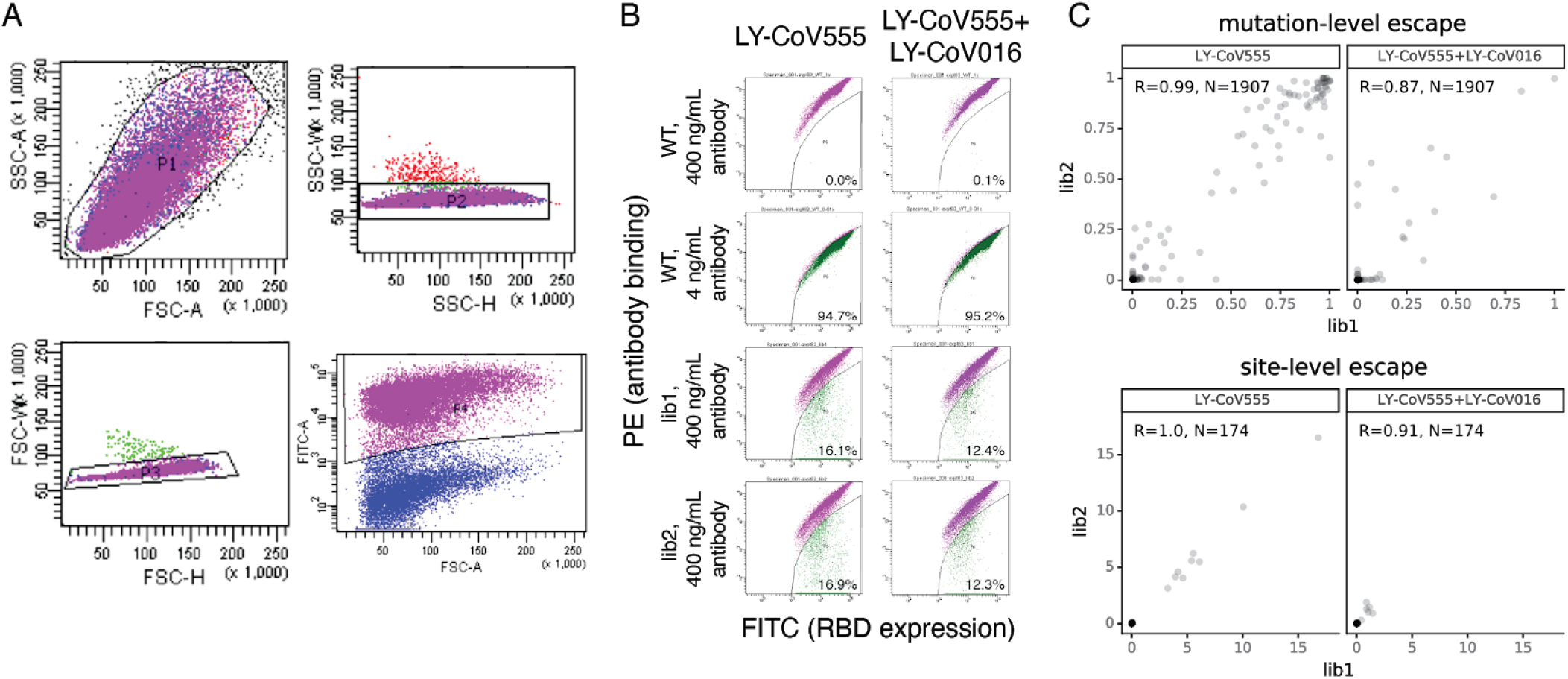
Experimental approach for antibody escape mapping. We use fluorescence-activated cell sorting to select yeast mutants that escape antibody binding. We use libraries described by Starr et al. [20] which contain virtually all RBD amino acid mutations, which were previously sorted to select for those compatible with human ACE2 binding as described by Greaney et al. [18]. (A) Initial FACS gates are drawn to select single yeast cells (SSC/FSC, SSC-W/SSC-H, and FSC-H/FSC-W) that properly express RBD (FITC/FSC). (B) Antibody-escape bins are drawn to capture 95% of RBD^+^ cells expressing the unmutated SARS-CoV-2 RBD labeled at 4 ng/mL, indicating 100x reduced binding compared to the 400 ng/mL library selections. The fraction of cells in each population that fall into this “antibody-escape” bin is labeled in each FACS plot. (C) Correlation between independent library duplicates at the level of per-mutation escape fractions (top) or site-level total escape (sum of all mutations at site, bottom).

**Supplemental Figure 2.**
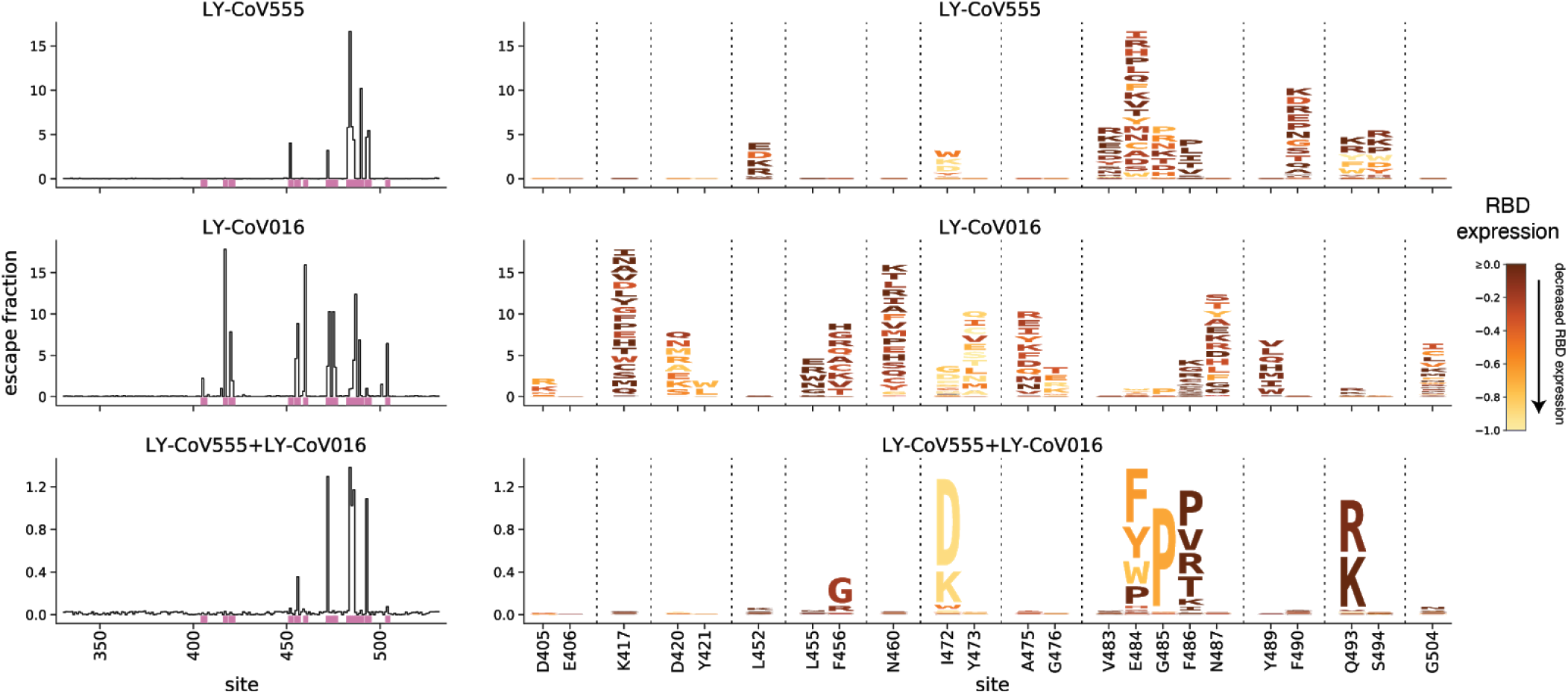
Logoplots colored by mutation effects on expression. Line and logoplots are identical to Fig. 1A, except mutations are colored by their effects on the yeast surface expression level of folded RBD, as measured in our previous deep mutational scanning experiment [20].

## Notes

https://jbloomlab.github.io/SARS-CoV-2-RBD_MAP_LY-CoV555/

### Competing Interest Statement

The authors have declared no competing interest.

## References

1. Chen P, Nirula A, Heller B, Gottlieb RL, Boscia J, Morris J, et al. SARS-CoV-2 Neutralizing Antibody LY-CoV555 in Outpatients with Covid-19. N Engl J Med. 2021;384: 229–237.

2. Weinreich DM, Sivapalasingam S, Norton T, Ali S, Gao H, Bhore R, et al. REGN-COV2, a Neutralizing Antibody Cocktail, in Outpatients with Covid-19. N Engl J Med. 2021;384: 238–251.

3. Tortorici MA, Beltramello M, Lempp FA, Pinto D, Dang HV, Rosen LE, et al. Ultrapotent human antibodies protect against SARS-CoV-2 challenge via multiple mechanisms. Science. 2020;370: 950–957.

4. Jones BE, Brown-Augsburger PL, Corbett KS, Westendorf K, Davies J, Cujec TP, et al. LY-CoV555, a rapidly isolated potent neutralizing antibody, provides protection in a non-human primate model of SARS-CoV-2 infection. bioRxiv. 2020. p. 2020.09.30.318972. doi:10.1101/2020.09.30.318972

5. Zost SJ, Gilchuk P, Case JB, Binshtein E, Chen RE, Nkolola JP, et al. Potently neutralizing and protective human antibodies against SARS-CoV-2. Nature. 2020;584: 443–449.

6. Hassan AO, Case JB, Winkler ES, Thackray LB, Kafai NM, Bailey AL, et al. A SARS-CoV-2 Infection Model in Mice Demonstrates Protection by Neutralizing Antibodies. Cell. 2020;182: 744–753.e4.

7. Rogers TF, Zhao F, Huang D, Beutler N, Burns A, He W-T, et al. Isolation of potent SARS-CoV-2 neutralizing antibodies and protection from disease in a small animal model. Science. 2020;369: 956–963.

8. Lilly’s neutralizing antibody bamlanivimab (LY-CoV555) receives FDA emergency use authorization for the treatment of recently diagnosed COVID-19. [cited 14 Feb 2021]. Available: https://investor.lilly.com/news-releases/news-release-details/lillys-neutralizing-antibody-bamlanivimab-ly-cov555-receives-fda

9. Lilly’s bamlanivimab (LY-CoV555) administered with etesevimab (LY-CoV016) receives FDA emergency use authorization for COVID-19. [cited 14 Feb 2021]. Available: https://investor.lilly.com/news-releases/news-release-details/lillys-bamlanivimab-ly-cov555-administered-etesevimab-ly-cov016

10. Tegally H, Wilkinson E, Giovanetti M, Iranzadeh A, Fonseca V, Giandhari J, et al. Emergence and rapid spread of a new severe acute respiratory syndrome-related coronavirus 2 (SARS-CoV-2) lineage with multiple spike mutations in South Africa. medRxiv; 2020. doi:10.1101/2020.12.21.20248640

11. Public Health England. Investigation of novel SARS-CoV-2 variant: Variant of Concern 202012/01. GOV.UK; 21 Dec 2020 [cited 11 Feb 2021]. Available: https://www.gov.uk/government/publications/investigation-of-novel-sars-cov-2-variant-variant-of-concern-20201201

12. Faria NR, Morales Claro I, Candido D, Moyses Franco LA, Andrade PS, Coletti TM, et al. Genomic characterisation of an emergent SARS-CoV-2 lineage in Manaus: preliminary findings. Virological; 2021. Available: https://virological.org/t/genomic-characterisation-of-an-emergent-sars-cov-2-lineage-in-manaus-preliminary-findings/586

13. Zhang W, Davis BD, Chen SS, Sincuir Martinez JM, Plummer JT, Vail E. Emergence of a Novel SARS-CoV-2 Variant in Southern California. JAMA. 2021. doi:10.1001/jama.2021.1612

14. Wang P, Liu L, Iketani S, Luo Y, Guo Y, Wang M, et al. Increased Resistance of SARS-CoV-2 Variants B.1.351 and B.1.1.7 to Antibody Neutralization. bioRxiv. 2021. p. 2021.01.25.428137. doi:10.1101/2021.01.25.428137

15. Starr TN, Greaney AJ, Addetia A, Hannon WW, Choudhary MC, Dingens AS, et al. Prospective mapping of viral mutations that escape antibodies used to treat COVID-19. Science. 2021 [cited 25 Jan 2021]. doi:10.1126/science.abf9302

16. Hoffmann M, Arora P, Gross R, Seidel A, Hoernich B, Hahn A, et al. SARS-CoV-2 variants B.1.351 and B.1.1.248: Escape from therapeutic antibodies and antibodies induced by infection and vaccination. bioRxiv. 2021. p. 2021.02.11.430787. doi:10.1101/2021.02.11.430787

17. Diamond M, Chen R, Xie X, Case J, Zhang X, VanBlargan L, et al. SARS-CoV-2 variants show resistance to neutralization by many monoclonal and serum-derived polyclonal antibodies. Research Square. 2021. doi:10.21203/rs.3.rs-228079/v1

18. Greaney AJ, Starr TN, Gilchuk P, Zost SJ, Binshtein E, Loes AN, et al. Complete Mapping of Mutations to the SARS-CoV-2 Spike Receptor-Binding Domain that Escape Antibody Recognition. Cell Host Microbe. 2021;29: 44–57.e9.

19. Greaney AJ, Loes AN, Crawford KHD, Starr TN, Malone KD, Chu HY, et al. Comprehensive mapping of mutations in the SARS-CoV-2 receptor-binding domain that affect recognition by polyclonal human plasma antibodies. Cell Host Microbe. 2021;0. doi:10.1016/j.chom.2021.02.003

20. Starr TN, Greaney AJ, Hilton SK, Ellis D, Crawford KHD, Dingens AS, et al. Deep Mutational Scanning of SARS-CoV-2 Receptor Binding Domain Reveals Constraints on Folding and ACE2 Binding. Cell. 2020;182: 1295–1310.e20.

21. Choi B, Choudhary MC, Regan J, Sparks JA, Padera RF, Qiu X, et al. Persistence and Evolution of SARS-CoV-2 in an Immunocompromised Host. N Engl J Med. 2020. doi:10.1056/NEJMc2031364

22. Shi R, Shan C, Duan X, Chen Z, Liu P, Song J, et al. A human neutralizing antibody targets the receptor-binding site of SARS-CoV-2. Nature. 2020;584: 120–124.

23. Raghuvamsi PV, Tulsian NK, Samsudin F, Qian X, Purushotorman K, Yue G, et al. SARS-CoV-2 S protein:ACE2 interaction reveals novel allosteric targets. Elife. 2021;10. doi:10.7554/eLife.63646

24. Guo H, Hu B-J, Yang X-L, Zeng L-P, Li B, Ouyang S, et al. Evolutionary Arms Race between Virus and Host Drives Genetic Diversity in Bat Severe Acute Respiratory Syndrome-Related Coronavirus Spike Genes. J Virol. 2020;94. doi:10.1128/JVI.00902-20

25. Shang J, Ye G, Shi K, Wan Y, Luo C, Aihara H, et al. Structural basis of receptor recognition by SARS-CoV-2. Nature. 2020;581: 221–224.

26. Eguia R, Crawford KHD, Stevens-Ayers T, Kelnhofer-Millevolte L, Greninger AL, Englund JA, et al. A human coronavirus evolves antigenically to escape antibody immunity. bioRxiv. 2020. p. 2020.12.17.423313. doi:10.1101/2020.12.17.423313

27. Kistler KE, Bedford T. Evidence for adaptive evolution in the receptor-binding domain of seasonal coronaviruses OC43 and 229e. Elife. 2021;10. doi:10.7554/eLife.64509

28. Weisblum Y, Schmidt F, Zhang F, DaSilva J, Poston D, Lorenzi JCC, et al. Escape from neutralizing antibodies by SARS-CoV-2 spike protein variants. Elife. 2020;9: e61312.

29. Andreano E, Piccini G, Licastro D, Casalino L, Johnson NV, Paciello I, et al. SARS-CoV-2 escape in vitro from a highly neutralizing COVID-19 convalescent plasma. bioRxiv. 2020. p. 2020.12.28.424451. doi:10.1101/2020.12.28.424451

30. Liu Z, VanBlargan LA, Bloyet L-M, Rothlauf PW, Chen RE, Stumpf S, et al. Identification of SARS-CoV-2 spike mutations that attenuate monoclonal and serum antibody neutralization. Cell Host Microbe. 2021. doi:10.1016/j.chom.2021.01.014

31. Otwinowski J, McCandlish DM, Plotkin JB. Inferring the shape of global epistasis. Proc Natl Acad Sci U S A. 2018;115: E7550–E7558.

32. Elbe S, Buckland-Merrett G. Data, disease and diplomacy: GISAID’s innovative contribution to global health. Glob Chall. 2017;1: 33–46.

33. Katoh K, Standley DM. MAFFT multiple sequence alignment software version 7: improvements in performance and usability. Mol Biol Evol. 2013;30: 772–780.

34. Lan J, Ge J, Yu J, Shan S, Zhou H, Fan S, et al. Structure of the SARS-CoV-2 spike receptor-binding domain bound to the ACE2 receptor. Nature. 2020;581: 215–220.

35. Hilton S, Huddleston$ J, Black A, North K, Dingens A, Bedford T, et al. dms-view: Interactive visualization tool for deep mutational scanning data. JOSS. 2020;5: 2353.

